# Highly accurate assembly polishing with DeepPolisher

**DOI:** 10.1101/2024.09.17.613505

**Authors:** Mira Mastoras, Mobin Asri, Lucas Brambrink, Prajna Hebbar, Alexey Kolesnikov, Daniel E. Cook, Maria Nattestad, Julian Lucas, Taylor S. Won, Pi-Chuan Chang, Andrew Carroll, Benedict Paten, Kishwar Shafin

## Abstract

Accurate genome assemblies are essential for biological research, but even the highest quality assemblies retain errors caused by the technologies used to construct them. Base-level errors are typically fixed with an additional polishing step that uses reads aligned to the draft assembly to identify necessary edits. However, current methods struggle to find a balance between over- and under-polishing. Here, we present an encoder-only transformer model for assembly polishing called DeepPolisher, which predicts corrections to the underlying sequence using Pacbio HiFi read alignments to a diploid assembly. Our pipeline introduces a method, PHARAOH (Phasing Reads in Areas Of Homozygosity), which uses ultra-long ONT data to ensure alignments are accurately phased and to correctly introduce heterozygous edits in falsely homozygous regions. We demonstrate that the DeepPolisher pipeline can reduce assembly errors by half, with a greater than 70% reduction in indel errors. We have applied our DeepPolisher-based pipeline to 180 assemblies from the next Human Pangenome Reference Consortium (HPRC) data release, producing an average predicted Quality Value (QV) improvement of 3.4 (54% error reduction) for the majority of the genome.

## Introduction

Faithful reconstruction of the genome is increasingly essential for scientists to understand the underlying biology of any organism. Genome assembly, the process of digitally reconstructing a genome, is accomplished by piecing together segments of DNA (called sequencing “reads”) and obtaining a consensus. The past several years have seen a huge improvement in the quality of genome assembly, driven by long-read sequencing technology, which produces reads that are orders of magnitude longer than the short-read technologies that preceded them^1,2^. Assembly algorithms have continued to advance alongside the improvements in sequencing, and it is now becoming routine to produce automated, fully-phased, highly contiguous, and near-complete genome assemblies^1–6^.

The quality of a genome assembly can be measured by an array of different metrics that assess different aspects of the reconstruction. Contiguity metrics describe the cumulative length and number of contigs of the assembly, while completeness metrics evaluate the extent of the expected sequence present in the assembly^1^. For organisms with multiple copies of their chromosomes (ploidy), fully resolving the haplotypes is a key component to accurate genome assembly, and chromosome phasing metrics assess this^7,8^. Another critical set of metrics evaluates the base-level accuracy of the assembled sequence^7^.

All sequencing technologies contain errors which are not completely random, but are influenced by biases towards different genomic features. Even using a consensus approach, recurrent errors in the reads can propagate into the assembly. Low complexity regions like homopolymers and tandem repeats are particularly challenging for Oxford Nanopore Technologies (ONT)^9^ and Pacific Biosciences (PacBio) long-reads^10^, and certain motifs such as those high in GC content can cause dropout, a phenomenon particularly prevalent in short-reads like Illumina^11,12^. Small errors can also be introduced by biases in the assembler, and different assembly algorithms differ in their limitations and error characteristics^4,5,13^.

Base-level errors interfere with scientists’ ability to make accurate conclusions from their data. For example, they can disrupt functional elements, as in protein coding genes, leading to frameshifts, missense, and nonsense mutations^14–16^. For projects like the Human Pangenome Reference Consortium (HPRC)^6^ and Vertebrate Genome Project (VGP)^17^, whose goal is to produce haplotype-resolved population-scale reference genomes for entire groups of species, fixing errors is especially critical to avoid propagating mistakes to all the downstream analyses that make use of them, for example, by proposing sequence variations that are in fact errors^16,18,19^.

The process of removing these small-scale errors in draft assemblies is known as “polishing”. Most methods for assembly polishing involve aligning reads back to the draft assembly to identify potential sequence changes suggested by the reads. Some methods employ heuristic algorithms, specialized models, or repurpose variant callers to identify polishing edits^20–22^. In order to polish CHM13 (an effectively haploid cell line), the T2T consortium leveraged DeepVariant^23^ calls from multiple sequencing technologies and a careful filtering strategy to avoid false positive polishing edits^24,25^. A major challenge for all polishing approaches is striking a balance between overcorrecting and under-correcting assemblies, due to the complex nature of sequencing and assembly error profiles.

Here we present a sequence-to-sequence (seq2seq) transformer-based method for assembly polishing. Seq2seq transformer family models have been used to great success in many applications including language processing, machine translation tasks, and conversational AI^26–28^. Previously this model architecture was used in DeepConsensus to improve the quality of PacBio High-Fidelity (HiFi) sequencing reads^29^. In this work, we demonstrate a new adaptation of the transformer model for genome polishing called DeepPolisher, which takes HiFi sequencing reads aligned to a draft assembly as input. Along with DeepPolisher, we have developed a pipeline called PHARAOH for improving the phasing accuracy of HiFi read alignments to diploid assemblies in falsely homozygous regions using ONT ultra-long (UL) reads. In this work, we show that DeepPolisher with PHARAOH outperforms current polishing approaches. Using an alignment-based measure of accuracy we demonstrate that DeepPolisher reduces assembly errors by half, driven mostly by reductions in assembly insertion-deletion (indel) errors. We applied our pipeline to 180 haplotype-resolved assemblies from the next HPRC data release, and show substantial improvements in accuracy, increasing the predicted quality value (QV) by 3.4 for most of the genome.

## Results

### Overview of the DeepPolisher pipeline

An overview of the DeepPolisher pipeline is shown in **Figure 1**. First, PacBio HiFi reads are aligned to the diploid assembly with minimap2^30^ or winnowmap^31^. For most of the genome, this alignment step is sufficient to assign the HiFi reads to the correct haplotype. However, a common error mode of modern assemblers is to mistakenly represent heterozygous sequence as homozygous. When these regions of false homozygosity are longer than the HiFi read (up to ~ 25kb), the aligner cannot phase it, and will randomly assign it to a haplotype, preventing any necessary polishing edits from being made there. To address this, we developed a pipeline called PHARAOH (PHAsing Reads in Areas Of Homozygosity) which uses phasing information from ONT reads greater than 100 kb to infer the correct haplotype for reads in long homozygous regions (**Methods**). For the rest of the genome, the tool Secphase^6^ is used within the PHARAOH pipeline to ensure correct read phasing by revisiting secondary alignments and calculating a marker consistency score to determine whether the read should be relocated to the other haplotype.

**Figure 1:**
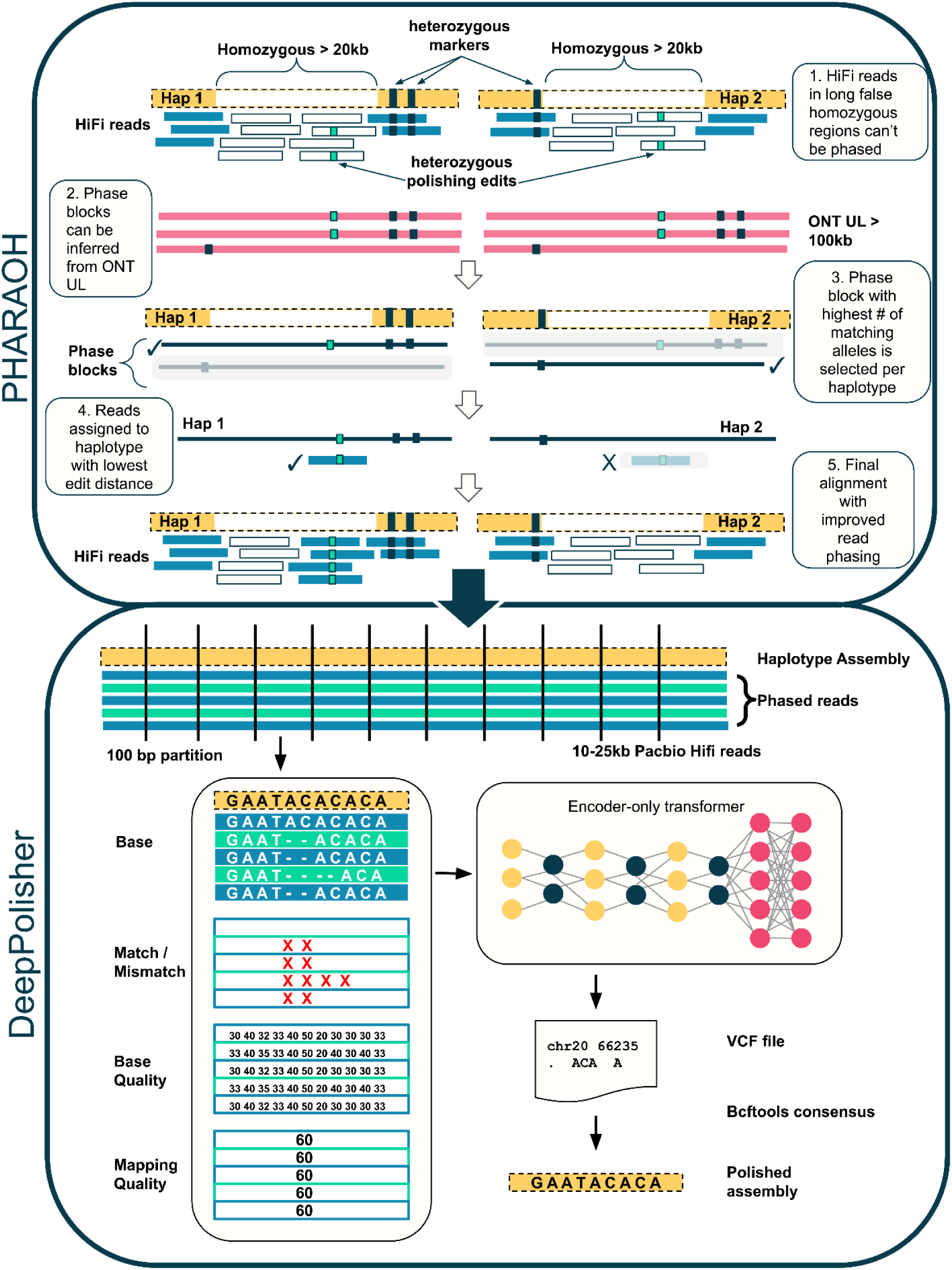
DeepPolisher pipeline overview. The PHARAOH pipeline leverages phase block information from ONT UL reads to correct the haplotype assignment of PacBio HiFi reads. The corrected alignment is passed to DeepPolisher, which is an encoder-only transformer model that predicts the underlying assembly sequence and proposes corrections in vcf format.

PHARAOH and Secphase allow us to provide improved read alignments as input to the DeepPolisher model. DeepPolisher first partitions these alignments into 100 bp segments, then represents them as a tensor object with auxiliary input features consisting of the base, whether the base is a match or mismatch to the assembly, the base quality, and the mapping quality. The tensor is provided to the encoder-only transformer neural network, which produces proposed assembly corrections in vcf format. These can be applied to the unpolished assembly with bcftools^32^ consensus (**Figure 1**). To train DeepPolisher we used a high accuracy assembly of HG002 produced by the Q100 consortium^33^ (v0.9), training across chromosomes 1-19 (**Methods**). DeepPolisher can be run without PHARAOH if ONT UL reads are not available, however assembly corrections in long stretches of false homozygosity will not be made.

### Alignment based comparison of DeepPolisher and alternate polishing approaches

We compared the performance of DeepPolisher with existing polishing approaches for HiFi reads: the Telomere-to-Telomere (T2T) consortium pipeline^24^, DeepVariant^23^ using HiFi alignments, and NextPolish2^20^. To test them, we ran the UL version of Hifiasm^4^ (which uses both HiFi and ONT UL reads) to generate an assembly for a sample that was not included in our training set (HG005). We polished it with each method, then called variants relative to GRCh38 using dipcall^34^, which enabled us to assess concordance with the Genome in a Bottle (GIAB) v4.2.1 benchmark call set^35^ (**Methods**). This gives an alignment-based measure of polishing accuracy that spans approximately 80% of the HG005 genome, as defined by the GIAB confident regions.

With DeepPolisher, we reduce the number of variant calling errors in the unpolished assembly from 20,274 to 11,750, indicating a reduction in assembly error rate by 43% (8.14 errors per Mb to 4.72 errors per Mb) **(Figure 2A)**. This performance is reproduced on the held-out chromosome (20) from HG002 (**Supplementary Table 1A,1B**). We also trained a DeepPolisher model on a Verkko^5^ assembly using the same data. According to the GIAB variant calling results, the Verkko assembly for HG005 starts out at a higher quality than Hifiasm (6.86 errors / Mb), but after DeepPolisher we bring it to the same quality as Hifiasm after polishing at 4.87 errors / Mb (**Supplementary Table 1E**).

**Figure 2:**
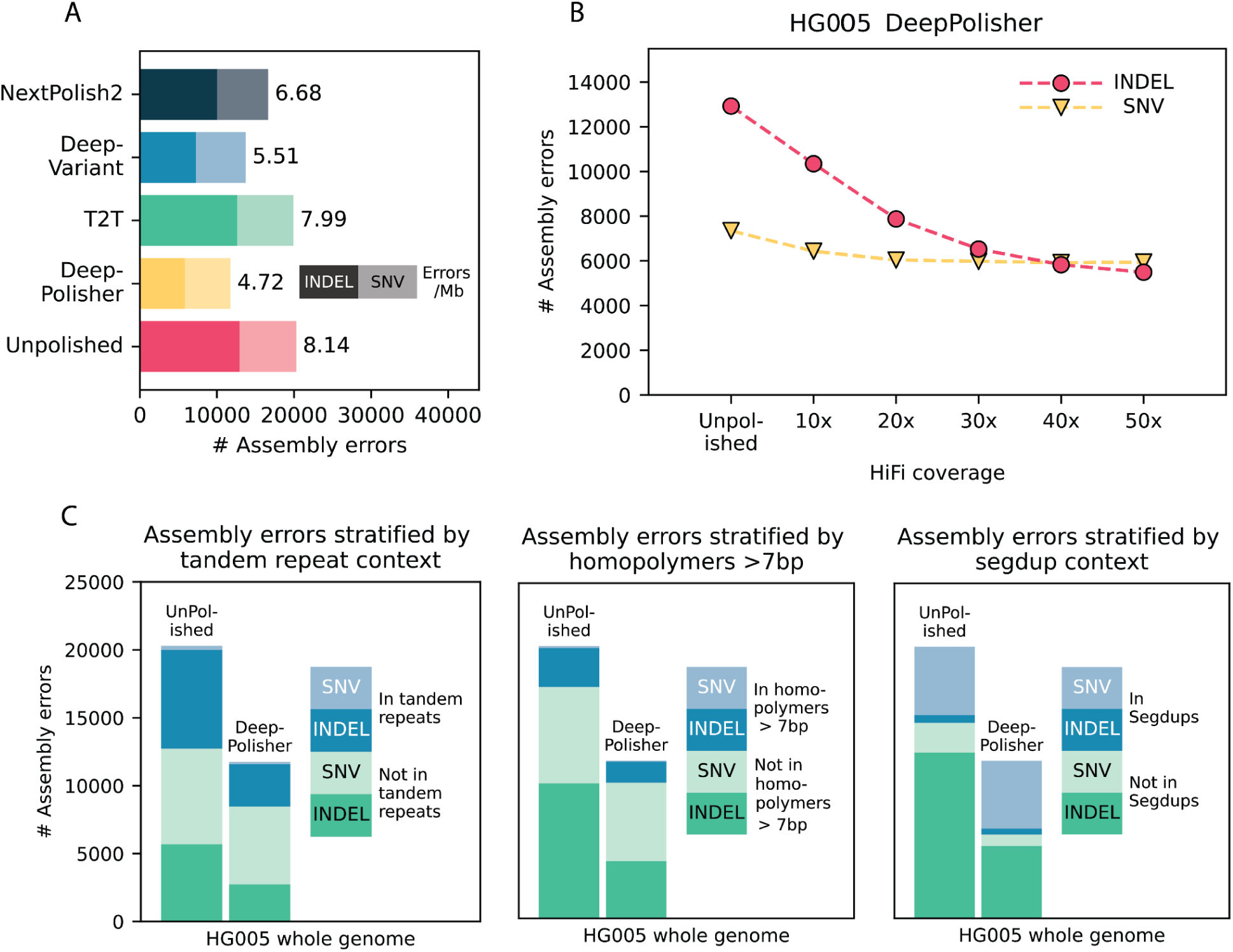
Comparison of DeepPolisher and alternate polishing methods against GIAB v4.2.1 benchmark for HG005. **A)** For each polishing method, GIAB v4.2.1 variant calling (assembly) errors are separated by indels (darker shade) and single nucleotide variants (SNVs) (lighter shade), with the number of errors per megabase to the right of each bar. **B)** Total GIAB variant calling (assembly) errors for different HiFi read coverages, with indel errors represented in pink circles and SNV errors in yellow triangles. **C)** Total GIAB variant calling (assembly) errors stratified by presence in tandem repeats (left), homopolymers > 7bp (middle) and segmental duplications (segdups) (right), with SNV errors in lighter shades and indel errors in darker shades.

In comparison to DeepPolisher, the T2T polishing pipeline produces more conservative improvements, with only 3,591 polishing edits passing their filters, leading to a removal of 372 errors (7.99 errors / Mb). Using DeepVariant to polish with HiFi alignments reduces assembly errors to 13,724 (5.51 errors / Mb). We note that much of the difference in GIAB variant calling performance between DeepVariant and DeepPolisher on HG005 is due to PHARAOH: polishing the HG005 assembly with DeepVariant run on PHARAOH alignments produces a reduction to 11,508 assembly errors (4.62 errors / Mb). However, an important caveat is that DeepVariant was trained on several GIAB samples including HG005, with only chr20-22 held out. Finally, NextPolish2 removed 3,260 errors from the unpolished assembly (6.68 errors / Mb) (**Figure 2A, Supplementary Table 1C**).

We tested DeepPolisher across a variety of coverages, and found that while 40x coverage produces optimal reduction in variant calling errors, some assembly improvement can still be obtained with as low as 10x coverage (**Figure 2B**). The DeepPolisher pipeline performance is also reproducible across HiFi read versions other than what it was trained on (DeepConsensus v1.2). This includes Sequel II, DeepConsensus v0.2, and Revio with DeepConsensus v1.1(**Supplementary Figures 1 and 2, Supplementary Table 2)**.

We stratified the GIAB variant calling errors by genomic region and found that the majority of indel errors removed by DeepPolisher were in tandem repeats and homopolymers. The majority of SNV errors that were not fixed by DeepPolisher were located within segmental duplications. Some of these may be true errors, but others may be mapping artifacts or mistakes in the truth set (**Figure 2C**).

To test how DeepPolisher impacts critical coding regions, we looked for nonsynonymous, frameshift, or premature stop mutations in the HG005 assembly before and after polishing. DeepPolisher removed two frameshifts, four nonsynonymous mutations, and one premature stop codon (**Supplementary Table 12**).

We ran DeepPolisher directly on minimap2 alignments to assess the performance benefits of the PHARAOH pipeline. DeepPolisher on PHARAOH alignments results in 1,915 fewer variant calling errors for HG002 (whole genome) and 519 less errors for HG005 (**Supplementary Table 1B, 1G)**.

### K-mer based comparison of DeepPolisher and alternate polishing methods

K-mer based methods assess the accuracy of an assembly using an alignment-free approach^7^. To compare the k-mer and alignment based assessments we first restricted the k-mer assessment to just the assembled sequence contained within the GIAB high confidence regions, which span ~80% of the HG005 assembly (**Methods**). K-mer methods generally report a quality value (QV), a log base 10 scaled measure of assembly error, with higher values indicating more accurate assemblies. We find that DeepPolisher improves the k-mer based assembly QV by 1.8, equivalent to a 34% reduction in errors, which is approximately consistent with the quantity of improvement suggested by the GIAB variant calling performance (**Figure 3A, Supplementary Table 6**). Out of all the polishing methods compared, DeepPolisher induces the least amount of error k-mers to the assembly (**Figure 3A**). NextPolish2, which uses Illumina k-mers in its algorithm for assembly polishing, produces the highest QV improvement, yet with the trade-off of inducing the highest number of new error k-mers to the assembly. The introduction of new error k-mers likely lies behind the increase in switch and hamming error rate produced by NextPolish2 (.03% and .06% respectively). In contrast, the other polishing methods manage to reduce switch and hamming error rates (**Figure 3B**).

**Figure 3:**
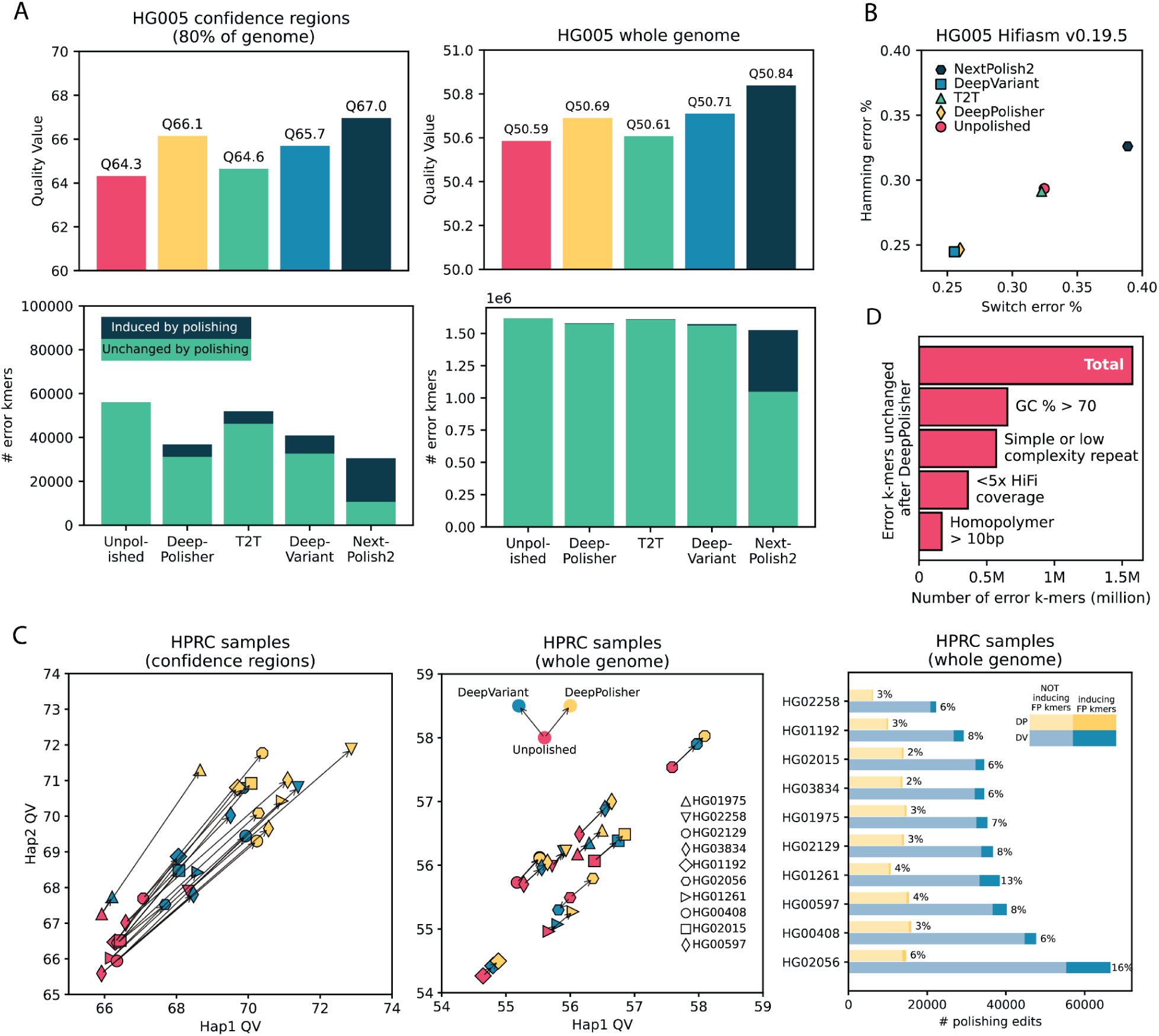
K-mer based comparison of DeepPolisher and alternate polishing approaches for HG005. **A)** Top panels display QV scores for each polishing method. Bottom panels depict total error k-mers, divided by error k-mers induced by polishing (dark blue) and error k-mers unchanged after polishing (green). Left panels show results for the GIAB confidence regions, right panels whole genome. **B)** Switch (x axis) and hamming (y axis) error rates for each polishing method. **C)** Comparison of DeepVariant and DeepPolisher for 8 HPRC samples. Left and middle panels show Hap1 (x axis) and Hap2 (y axis) QV for 8 HPRC samples, with an arrow connecting the unpolished QV (pink) to the QV after polishing with DeepVariant (blue) and DeepPolisher (yellow). Left panel is within the GIAB confidence regions, middle panel whole genome. Right panel shows number of polishing edits from DeepPolisher (yellow) and DeepVariant (blue). Lighter shades indicate edits not inducing error (FP) k-mers, darker shades show edits that induce error k-mers. **D)** Number of error k-mers unchanged by polishing with DeepPolisher falling into sequence annotation categories.

To compare the performance of DeepVariant and DeepPolisher on samples that were completely held out from training in both methods, we applied both tools to the same PHARAOH alignments for eight of the HPRC samples, and assessed their QV. For all samples, DeepPolisher produces a higher QV improvement (avg. 3.95 QV, 60% error reduction) within the confidence regions compared with DeepVariant (avg 2.26 QV, 41% error reduction) (**Figure 3C**). In addition, the percentage of edits inducing error k-mers is more than twice as high for DeepVariant (average 9%) than DeepPolisher (average 4%) (**Figure 3C**).

### K-mer based assessments underestimate assembly error rates

Relative to alignment-based metrics like those used in the GIAB assessment above, k-mer-based QVs are unbiased by alignment and easy to assess. However, k-mer based methods lack an underlying ground truth, instead relying on consistency between assembly and sequencing technology. This results in a potential limitation where results could be consistent but not correct due to technology-specific sequencing bias. Because k-mer based quality metrics are agnostic to the sequence context outside of the k-mer window, they also are limited in their ability to capture errors in repetitive sequence or in regions with a loss of heterozygosity.

For the unpolished HG005 assembly within the confident regions, we see a QV estimate of 66.97 using Illumina k-mers (k-31), suggesting 0.811 errors / Mb, while the GIAB analysis suggests the same assembly has ~10x more errors per Mb (8.14) (equivalent to a QV of 50.9) **(Supplementary Table 6)**. Using shorter k-mer sizes or a hybrid k-mer set built with both Illumina and HiFi reads as suggested by McCartney et. al^24^ led to substantially more inflated estimates of accuracy (0.258 and 0.201 errors / Mb, respectively) (**Methods, Supplementary Table 6)**. In order to explore the source of this difference in error rate, we performed a manual analysis investigating the concordance between the errors reported by the QV and the GIAB variants for HG005 after polishing with DeepPolisher (**Methods, Supplementary Table 13**). We found that error k-mers reported by QV were almost always validated by errors reported using GIAB, indicating that most of the QV errors within the confident regions are likely true. However, for 35% of the cases examined, the QV did not flag the pre- nor post-polishing sequence as an error. Because it is impossible for both versions of the assembly sequence to be correct, we attribute this to the fact that k-mer based quality estimates often miss errors in repetitive sequences, due to their presence in the “truth” k-mer set coming from other regions of the genome. Therefore, we conclude that non-unique sequence is likely the cause of the underestimated error rates in QV metrics.

### K-mer based methods identify residual errors in unpolished sequence

Looking at the k-mer QV whole genome we found little difference between the unpolished assembly and the four different polishing methods, with all QV scores in the range of 50.56 - 50.84. This is because the vast majority of the error k-mers leading to this QV score are unchanged after polishing (97.3% for DeepPolisher) (**Figure 3A**). Investigating these unchanged error k-mers, we found that 41% have greater than 70% GC content, and when we plotted the distribution of their GC content we saw a clear bias relative to randomly permuted k-mers (**Methods**, **Supplementary Figure 4**). This indicates that the observed GC bias^11,12^ in short reads is likely inducing a substantial number of falsely reported errors. However, this still leaves a majority of these error k-mers that may be real, unaddressed errors. Indeed, we find that 22.8% of predicted error k-mers are located in regions with less than 5x coverage of HiFi data. These regions were likely bridged by ONT sequence in the assembly stage, and therefore have a higher error rate, but are inaccessible to DeepPolisher due to the lack of HiFi read coverage. Additionally, we found that 36% of the unchanged error k-mers overlapped simple and low-complexity repeat annotations, with 10% overlapping homopolymers greater than 10 bp. These regions tend to have higher error rates in HiFi reads^10,36^, which makes it likely these are true errors that DeepPolisher was unable to fix. Together, these observations indicate that while the residual predicted error k-mers contain many false positives, the k-mer based methods are identifying significant sequence outside of GIAB high-confident regions that are erroneous and unchanged by our current polishing methods (**Figure 3D**, **Supplementary Table 7**).

### Polishing the HPRC release 2 assemblies

We applied our polishing pipeline to 180 HPRC assemblies made available as part of the second HPRC data release. Within the confident regions (81.4% of the genome on average), and consistent with our analysis of HG005, we noted an average QV improvement of 3.4 (54% reduction in errors), with the average unpolished QV of 66.66 moving to an average of 70.05 after polishing (**Figure 4A**). Whole genome, we improved the QV by 0.306 points on average (**Figure 4B**; 7% reduction in errors), owing to the presence of the lower quality ONT-only regions that cannot be polished with the current model, and regions subject to sequencing bias as previously discussed. Polishing improved phasing accuracy for all 111 samples that contained trio data: on average it reduced switch error by 0.039 and hamming errors by 0.037 **(Figure 4C**).

**Figure 4:**
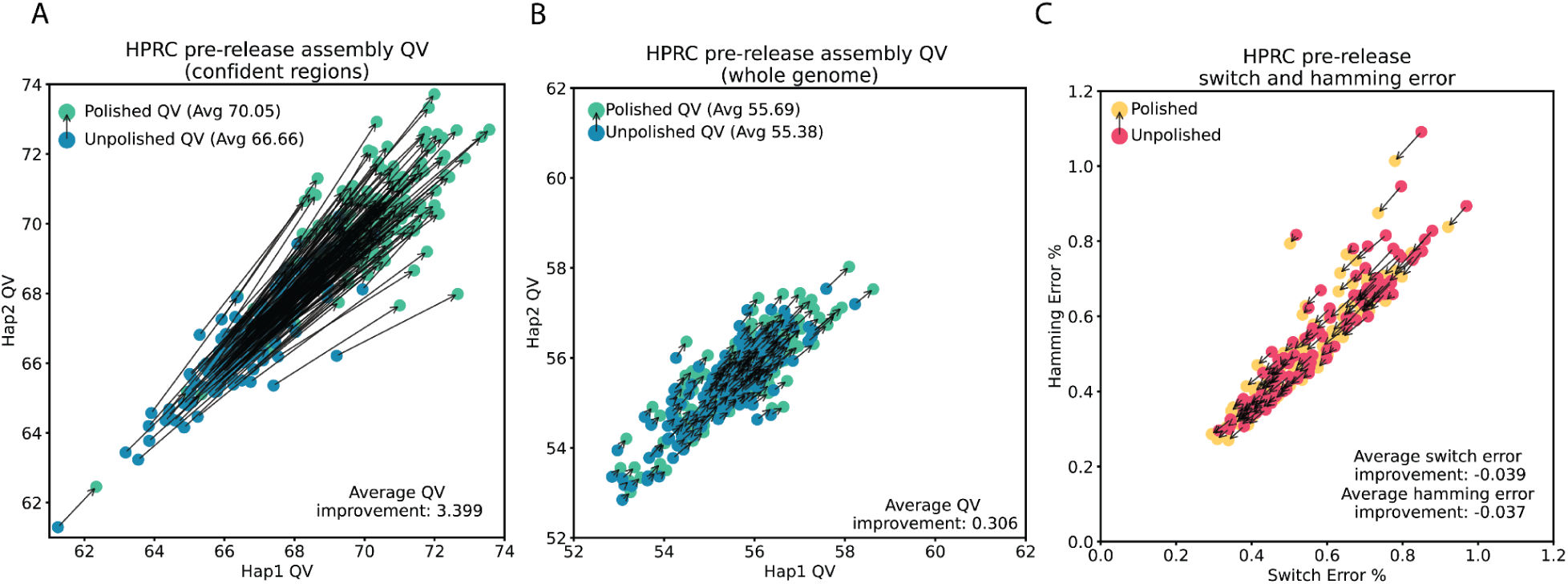
Polishing results for 180 HPRC assemblies. **A)** Hap1 QV (x axis) and Hap2 QV (y axis) in the confidence regions for 180 HPRC samples from the second release. For each sample, unpolished QV is in blue with an arrow pointing to the polished QV. **B)** The same as A) but for whole genome QV. **C)** Switch (x axis) and hamming (y axis) error rate for the 107 samples with trio data. Unpolished in pink with an arrow pointing to polished in yellow.

### Removing additional errors with Element data

Element Biosciences Avidity short-read data has gained popularity recently for its high accuracy in tandem repeats and homopolymers. We tested whether further polishing with Element data after DeepPolisher could fix residual errors. Because of the challenges associated with phasing short read data, we restricted our initial experiment to polishing just homozygous locations. In order to obtain potential homozygous edits, we aligned 50x Element cloudbreak reads^37^ to each haplotype of the HG002 and HG005 polished assemblies. We ran DeepVariant on each alignment, and selected just the homozygous-alternate calls with a GQ greater than 7. This approach removed 9% (1,182) of the remaining variant calling errors after polishing with DeepPolisher for HG002, and 5% (600) for HG005, a promising number of errors removed given the relatively low number of polishing edits applied (5,586 for HG002 and 8,795 for HG005) (**Supplementary Table 11**).

## Discussion

Rapid advancements in long-read sequencing and assembly algorithms in recent years are ushering in a new era of population-scale, T2T genome assembly projects^6,17,38^. Many of these assemblies are being used as reference genomes, making it important that the sequences be highly accurate. To remove the remaining errors in these sequences, deep learning approaches are a promising way to model sequencing bias in regions like homopolymers and low-complexity repeats. In this work, we developed a new polishing tool called DeepPolisher, an encoder-only transformer model, that takes in HiFi reads aligned to a diploid assembly, and predicts the underlying sequence. Accompanying this, we introduced a pipeline called PHARAOH (PHAsing Reads in Areas of Homozygosity) that ensures read alignment inputs to DeepPolisher are assigned to the correct haplotype, and phases potential polishing edits in long homozygous assembly regions.

Using an alignment based assessment of assembly accuracy measured against the GIAB v4.2.1 benchmarking variant calls, we demonstrate that DeepPolisher reduces assembly errors by approximately half, largely driven by reductions in indel errors, across the large majority of sequence that is mappable by HiFi sequencing. While alignment-based assessments of high-quality benchmark sets are close to a gold standard, current benchmarks do not cover the full genome and are dependent on accurate alignment. To get around these issues we also assessed assembly quality with k-mer methods. Comparing k-mer methods to the alignment-based assessment within the same genomic regions predicted similar percentage reductions in errors, which is consistent and reassuring, but also revealed that the k-mer methods miss around 90% or more of residual errors. While some of these may be false positives in the alignment benchmark, our manual analysis suggests that most are simply missed by the k-mer methods due to their limitations regarding modeling repeat k-mers that are likely often enriched in assembly errors. We would therefore urge users to recognize this and note that such QV estimates are likely overly optimistic because they have a high false negative rate.

However, k-mer methods also reveal large numbers of predicted errors in regions that remain untouched with the current iteration of DeepPolisher and are located outside of the regions assessed by the alignment based benchmarks. While nearly half of these predicted errors may be false positives caused by GC content bias, it is probable that many are real. We attribute a quarter of these errors to regions of HiFi coverage dropout where ONT was primarily used during the assembly stage. We plan to expand the DeepPolisher model to work on ONT reads in the future, to address this fraction of errors. Around a third of the residual errors lie in homopolymers and tandem repeats, sequences that are challenging even for highly accurate HiFi reads. We experimented with using Element Biosciences data to fix these errors and demonstrate that we can remove a minority of errors using the approach. However, Element reads are currently relatively short, preventing accurate mapping across repetitive sequence and limiting phasing. We predict the remaining class of errors will be challenging to fix.

To resolve the limitations of current alignment and k-mer based approaches to assessing base-level assembly accuracy, the Q100 project^33^ aims to create a “perfect” genome assembly to be used as a benchmark for assembly methods. While achieving a completely perfect assembly will be challenging, the direct comparison of a draft assembly to an assembly benchmark will help expand the alignment-based assessment of assembly quality to more difficult regions of the genome than are covered by the GIAB confidence regions. In the future, we hope that genome benchmarks for more samples will be produced, to expand the training and testing sets to multiple genomes for models like DeepPolisher.

Despite the difficulties inherent in assessing the performance benefits of the DeepPolisher model, we demonstrate that it provides the best balance between over and under-polishing for high-quality assembly projects like the HPRC. We have applied the DeepPolisher and PHARAOH pipeline to the next release of HPRC assemblies, and demonstrate consistent improvements to quality value across all samples, which will be critical for downstream applications of this new reference dataset. Next, we aim to experiment with DeepPolisher on non-human species, to expand its utility to high quality genomes outside of humans, which may be useful for projects like the VGP^17^ and T2T primates consortium^39^.

## Code availability

DeepPolisher code is available publicly on GitHub through (https://github.com/google/deeppolisher). PHARAOH is implemented in a pipeline available on GitHub (https://github.com/miramastoras/PHARAOH) Scripts for analysis and benchmarking are available on GitHub (https://github.com/miramastoras/DeepPolisher_manuscript). Code used to generate the HPRC assembly data release 2 may be found at (https://github.com/human-pangenomics/hprc_intermediate_assembly)

## Data availability

All data used to generate and polish the assemblies was downloaded from the HPRC at (https://s3-us-west-2.amazonaws.com/human-pangenomics/index.html?prefix=working/HPRC_PLUS/). A list of the specific files used in each step may be found at (https://github.com/miramastoras/DeepPolisher_manuscript/blob/main/assembly/hifiasm_HG002_HG005.md). The HPRC release 2 assemblies and their associated data may be found at (https://github.com/human-pangenomics/hprc_intermediate_assembly/tree/main/data_tables)

## Acknowledgements

B.P. and UCSC personnel were in part supported by NIH grants R01HG010485, U01HG010961, U24HG010262, OT2OD026682, and U24HG011853. M.M was in part supported by NIH grant T32HG012344.

## Author contributions

K.S. and B.P. devised the study. M.M., B.P., and K.S. drafted the manuscript. A.C, D.E.C, P.C., A.K., L.B., M.N. and K.S. developed DeepPolisher. M.A. and M.M. developed PHARAOH. M.M., M.A., and T.S.W. performed data analysis. P.H. performed gene impact analysis. M.A. and J.L. generated assemblies.

## Methods

### Assembly

To create HG002 and HG005 Hifiasm assemblies for training and testing DeepPolisher, we ran Hifiasm (UL) v0.19.5 in trio mode with default parameters. To create HG002 and HG005 Verkko assemblies for training and testing DeepPolisher, we ran Verkko v2.0 in trio mode with --correct-overlap-batches 64 and --screen human and otherwise default parameters. As input we used 40x HiFi DeepConsensus v1.2 and 40x ONT UL >100kb reads from the HPRC. The ONT reads for the HG005 and HG002 samples were sequenced using R9.4.1 flow cells and basecalled using Guppy software (HG005 with version 5.0.7 and HG002 with version 6.0.6). For phasing assembly haplotypes in the trio mode, both assemblers needed k-mer databases that were created from parental short reads, and Verkko required k-mer databases from the child’s short reads. For Hifiasm we used yak and for Verkko we used meryl to create the related k-mer databases. For HG002, we used the 300x parental and child illumina data from the HPRC to create input yak and meryl databases (using default settings for both), and 100x illumina data for HG005 parental and child databases.

### DeepPolisher development and training

DeepPolisher uses an encoder-only transformer model to predict potential errors present in a haplotype-resolved genome assembly. The framework of DeepPolisher is adopted from our previous work DeepConsensus^29^. The input to DeepPolisher is an alignment file where the reads are aligned to the correct haplotype assembly. Then it performs the following steps to identify potential errors:

1. **make_images**: In the make_images step:

○ DeepPolisher takes a 25kb window of the assembly and tries to find potential positions where there could be an error. This is done by comparing the reads to the assembly. If at any position, there are more than two reads containing a sequence that does not match the assembly (mismatched base, insertion or deletion), we pick that position to be a potential error site.
○ The positions that are within 50 bp or closer are grouped together to create a window that does not exceed 100 bp sequence.
○ For each window, we take alignment features (base, base quality, mapping quality, match or mismatch) and create a tensor-like representation of the window that potentially contains assembly errors.
○ This tensor-like representation of the alignment is then fed to the transformer-model for prediction.
2. **inference**: In the inference step, we take the tensor-like representation of the 100 bp window with potential sequencing errors and use that as an input to the encoder-only transformer model. The model predicts a sequence based on the read alignments. We then compare the predicted sequence to the assembly and any difference between the predicted sequence and the assembly is reported in a vcf format file which would indicate potential errors in the assembly. These vcf edits can be applied to the assembly after DeepPolisher in a separate step using bcftools^32^ consensus -H2.

We trained DeepPolisher on the HG002 Hifiasm assembly taking the HG002-T2T-v0.9^33^ as the truth. We took HiFi reads aligned to the Hifiasm assembly using the PHARAOH pipeline. Then we aligned the Hifiasm diploid assembly to HG002-T2T-v0.9 assembly to create an assembly-to-truth alignment, which we use as the truth sequence for training the model. During training, the predicted sequence is compared against the truth sequence and the loss is calculated via the alignment loss function we introduced in our previous work DeepConsensus^29^. We use a similar approach to train a model for Verkko assembly, where we use the Verkko assembly for training.

### PHARAOH pipeline

The first step in the PHARAOH workflow involves aligning all HiFi reads to the diploid assembly using minimap2 with parameters “-L --eqx --cs -c -k19 -x map-hifi”. Next, we identify stretches of homozygosity in the assembly by aligning the two haplotype fasta files to each other with minimap2 using the parameters “-L --eqx --cs -c -x asm5”. The resulting paf file is passed into a python script (https://github.com/mobinasri/secphase/blob/main/programs/src/find_homozygous_regions.py) which parses the cigar string to return stretches of 100% identical sequence greater than 20,000 bp (the average length of HiFi reads). Next, the reads within the homozygous regions are extracted and aligned separately to the maternal and paternal haplotypes using minimap2 with parameters “-L --eqx --cs -c -k19 -map-hifi”. All reads with gap-compressed mismatch ratio exceeding 0.02 are removed to avoid calling spurious variants in reads too diverged from the assembly. DeepVariant is then used to detect heterozygous variants in these alignments, and all variants with a genotype quality less than 10 are filtered out. We align ONT UL reads greater than 100,000 bp separately to each haplotype assembly using minimap2 with parameters “--cs --eqx -L -Y -map-ont”. These UL alignments are passed to WhatsHap^40^, which is used to phase the heterozygous variants called by DeepVariant in homozygous regions.

In order to reassign reads to the correct haplotype assembly using the phased variants from WhatsHap, we added a new mode to the tool Secphase. From WhatsHap we have phased variants grouped into phase blocks. Each phase block contains two haplotypes, and Secphase picks the haplotype with a higher number of reference alleles (in other words, the one that is more similar to the assembly sequence) covered by that phase block. Secphase then retains the variants with alternative alleles in the selected haplotype of the phase block. Next, a variant block is created around each selected variant. The minimum block size can be set as a parameter (default = 100) and the actual size is adjusted to be twice the variant size if the minimum block size is smaller than twice the variant size. Overlapping variant blocks are merged, so a single block may encompass multiple variants. Each variant block, defined in assembly coordinates, is then projected onto the read coordinates. A single read may have multiple alignments, typically one per assembly haplotype, with each alignment having its own projected blocks. To create non-overlapping blocks crucial for distinguishing assembly haplotypes, all projected variant blocks are merged on the read coordinates. These merged blocks serve as proxies to compare haplotypes. The variants are then applied to the assembly haplotypes, effectively “pseudo-polishing” the assembly only in the created blocks. The pseudo-polished sequence of each haplotype is then compared against the read sequence within each variant block, with edit distances calculated using the Edlib^41^ library. The edit distances across all merged variant blocks are summed for each alignment. The haplotype with the lower total edit distance is selected as the correct haplotype for the related read.

In the PHARAOH pipeline, this new variant mode of Secphase is used for the homozygous regions > 20 kb containing a potential heterozygous polishing edit, and the original marker mode is used for the rest of the genome. After reassigning reads to the new location determined by Secphase, we remove alignments with a gap-compressed mismatch ratio exceeding 0.002, producing the final alignment for input to DeepPolisher. PHARAOH is implemented as a wdl workflow, and is publically available at https://github.com/miramastoras/PHARAOH. The tool Secphase may be found at https://github.com/mobinasri/Secphase. The entire pipeline for running PHARAOH and DeepPolisher may be found at https://github.com/human-pangenomics/hpp_production_workflows/tree/master/polishing.

### DeepPolisher GQ filter optimization

We selected four HPRC samples (HG01975, HG04115, HG02129, HG01993) with varying QV improvements after polishing (**Supplementary Table 10**) to optimize a set of GQ filters for DeepPolisher. First, we annotated the error k-mers in the assemblies by whether they were induced, fixed or didn’t change with polishing. To accomplish this we first aligned each haplotype of the unpolished assembly to the corresponding haplotype of the polished assembly with minimap2 and parameters -x asm5 -L --eqx --cs -c. We took the polished assembly *_only.bed file produced by Merqury (which contains the assembly coordinates for the error k-mers), merged the k-mers with bedtools^42^ merge -c 1, and projected them to the unpolished assembly using the script https://github.com/mobinasri/flagger/blob/main/programs/src/project_blocks_multi_thread.py. This allowed us to have the polished assembly error k-mers in unpolished assembly coordinates. We then subtracted the polished projected error k-mer bed file from the raw error k-mer bed file with bedtools subtract -A to extract the error k-mers fixed by polishing. We performed the reverse to obtain the error k-mers induced by polishing. Finally, we used bedtools intersect to produce the error k-mers unchanged by polishing, which were common to both bed files. To annotate the polishing edits by whether they induced, fixed, or didn’t change error k-mers, we intersected the polishing vcf with the error k-mer bed files labeled in the previous step.

We found that the majority of DeepPolisher edits inducing error k-mers were 1bp insertions or deletions (**Supplementary Figure 4**). Based on this observation, we tested three filtering scenarios: 1) A single GQ filter for all variants, 2) a GQ filter for 1bp insertions and a separate one for the remaining edits, and 3) a separate GQ filter for 1bp insertions, 1bp deletions, and for all other edits. We combined the annotated polishing edits from all four samples, and for each of these scenarios, we swept the GQ filter cutoff from 0 to 25 and calculated the estimated QV improvement using the formula -10*log10(error_k-mer_after/error_k-mer_before). We found the optimal GQ cutoff to be GQ 20 for 1 bp insertions, GQ 12 for 1 bp deletions, and GQ 5 for all other variants (**Supplementary table 10**). All DeepPolisher results shown in the paper have this filter applied unless otherwise stated. Detailed code for this analysis may be found at https://github.com/miramastoras/DeepPolisher_manuscript/blob/main/paper_analysis/Optimizing_GQ_filters.md.

### Implementing alternate polishing methods

We adapted the T2T polishing pipeline described in McCartney et. al^24^ for diploid assemblies to compare against DeepPolisher. We aligned 40x HiFi DeepConsensus v1.2 reads from the HPRC for both cell lines to the diploid Hifiasm assemblies with winnowmap v2.03 and parameters --cs --eqx -L -Y -I8g -map-pb. We aligned ONT R9 40x reads from the HPRC to the diploid Hifiasm assembly using winnowmap v2.03 and parameters --cs --eqx -I8g -Y -L -p0.5 -map-hifi. We also aligned 30x Illumina data to the diploid Hifiasm assembly with bwa-mem v.0.7.17 and default parameters. HiFi and Illumina bam files were merged with samtools, and DeepVariant v1.6.1 with parameter –model-type=HYBRID_PACBIO_ILLUMINA was used to call variants. PEPPER-Margin-DeepVariant was used to call variants for the ONT alignments. All GQ and VAF filters described in McCartney et al^24^. were applied. Merfin mode-polish was also used to further filter polishing edits. Edits were applied to the assembly with bcftools consensus -H1.

We used the same 40x HiFi DeepConsensus v1.2 winnowmap read alignments described above as input to NextPolish2 and DeepVariant. For NextPolish2, 30x Illumina data from the HPRC was first pre-processed with the tool fastp^43^ with parameters -f 5 --cut_front --cut_tail as suggested by the authors. The preprocessed data was then used to create a yak k=21 and k=31 database as input to NextPolish2, using default settings. NextPolish2 was run with default settings. DeepVariant v1.6.1 was run with default settings and flag --model_type=PACBIO. For DeepVariant, bcftools consensus -H1 was used to apply edits to the original assembly. All code used to implement alternate polishing methods is available at https://github.com/miramastoras/DeepPolisher_manuscript/blob/main/paper_analysis/alternate_polishing_methods.md

### Measuring GIAB variant calling performance

To assess the accuracy of the assemblies against the GIAB v4.2.1 benchmark set, we first ran dipcall^34^ with default settings against GRCh38 obtained from (https://ftp-trace.ncbi.nlm.nih.gov/ReferenceSamples/giab/release/references/GRCh38/). We next ran the tool hap.py^44^ to calculate performance metrics of the dipcall vcf file against the GIAB v4.2.1 benchmark vcf.

For HG002, we restricted the hap.py analysis to regions within the GIAB high confidence subset that were also validated by the T2T Q100 v1.0.1^33^ assembly. To create this subset, we ran dipcall^34^ on the T2T v1.0.1 assembly against GRCh38 using minimap parameters -z200000,10000 (in order to align across larger SVs and more divergent regions like the MHC). We next intersected the HG002 high confidence bed file provided by GIAB with the bed file produced by dipcall in the previous step, to produce a bed file of high confidence regions that are alignable between GRCh38 and T2T v1.0.1. We ran hap.py on the output vcf of dipcall vcf containing variants between T2T v1.0.1 and GRCh38 against the GIAB v4.2.1 vcf, restricting the analysis to the high confidence regions that are alignable between GRCh38 and T2T v1.0.1 by passing that bed file in with parameter -f. Using bedtools subtract we removed 50 bp surrounding any FP or FN designated variants produced in that hap.py run from the GIAB high confidence bed file, in order to remove regions that were not concordant with the T2T v1.0.1. 830 total variants were subtracted, and 63.7% of them were located within the GRCh38 segmental duplications track. Code for this analysis may be found at (https://github.com/miramastoras/DeepPolisher_manuscript/blob/main/paper_analysis/GIAB_truthset.md).

To create the final bed file to use with hap.py -f in benchmarking the HG002 assemblies, we intersected the bed file containing regions concordant between the Q100 T2T v1.0.1 assembly and GIAB v4.2.1 (described in the previous paragraph) with the bed file produced from the dipcall output of the unpolished assembly to GRCh38. For HG005, we repeat this step but with the original HG005 high confidence bed file provided by GIAB. For all hap.py runs we used the options --pass-only --no-roc --no-json --engine=vcfeval, and the GIAB v3.3 stratifications to obtain performance statistics per genomic region.

### Stratifying QV by whole genome and within GIAB confidence regions

For QV calculations we ran Merqury and Yak with default settings, using ~30x coverage Illumina data for all samples. For trio samples, we used Yak trioeval along with parental Illumina data to calculate switch and hamming rates. In order to assess QV just within the confidence regions, we first took an intersection of the GIAB high confidence regions for HG002 and HG005 using bedtools to define a subset of high confidence regions that isn’t genome-specific. We aligned each haplotype assembly to GRCh38 with minimap2 and parameters -x asm5 -L --eqx --cs -c. We then used the script https://github.com/mobinasri/flagger/blob/main/programs/src/project_blocks_multi_thread.py to lift over the GIAB confidence regions bedfile from GRCh38 to the assembly. Next, we used bedtools getfasta with default settings to extract only the sequence found within this bed file, and passed it to Merqury and Yak for QV calculation. In order to count the number of error k-mers induced, fixed, or unchanged after polishing, and obtain the polishing edits which were the cause of inducing, fixing, or failing to change the error k-mers, we performed the same annotation procedure described under the section “DeepPolisher GQ filter optimization**”.** To transform QV value into an estimate of error per megabase, we used the formula (10^(−QV / 10))* 10^6). This procedure is implemented in a wdl workflow publicly available at https://github.com/miramastoras/DeepPolisher_manuscript/blob/main/wdl/workflows/hprc_polishing_QC.wdl.

### Manual investigation of concordance between GIAB variant calls and Illumina QV

We performed a manual investigation of the concordance between the GIAB variant call errors and the Illumina QV reported errors for HG005 (polished by DeepPolisher) within the high confidence regions using IGV^45^. In order to obtain bed tracks for viewing the GIAB variant call errors in respect to the raw assembly coordinates, we used bcftools^32^ isec to obtain the intersections, unions and complements between the raw and polished vcfs output by hap.py (these vcfs contain variants relative to GRCh38 labelled by true-positive, false-positive or false-negative). We used the tool vcf2bed from BedOps^46^ to convert the vcf files to bed format, expanded the intervals by 10 base pairs on each side to enable projecting of the indels over to the raw assembly coordinates with the script https://github.com/mobinasri/flagger/blob/main/programs/src/project_blocks_multi_thread.py. For viewing the QV errors in IGV, we took the *_error.bed file (containing the locations in the assembly of the error k-mers) produced by Merqury using Illumina k-mers of size 31, and used the procedure described in “DeepPolisher GQ filter optimization**”** to annotate the polishing vcf by whether the edits induced, fixed, or failed to change the error k-mers, as well as label the error k-mers by whether they were fixed, induced or unchanged by polishing.

We used the files described above to randomly select 10 GIAB errors induced by DeepPolisher, 10 GIAB errors fixed by DeepPolisher, 10 polishing edits inducing error k-mers, and 10 polishing edits removing error k-mers. We carefully evaluated all 40 examples in IGV alongside the PHARAOH read alignments. For the GIAB errors we looked at both haplotypes to understand if one or both alleles was considered an error, and in two cases there was a polishing edit made to both haplotypes for a given GIAB error, so edits on both haplotypes were assessed. For all 42 polishing edits, we recorded the error status in both GIAB and QV. All details for the 42 edits may be found in **Supplementary Table 13**.

In summary, out of 42 polishing edits manually inspected, there were 3 cases where GIAB indicated the edit induced an error while the QV indicated that it fixed an error. For 24 / 42 polishing edits, the QV and GIAB metrics both agreed on whether the edit fixed or induced an error. For 15 / 42 polishing edits, the QV reported no error k-mer in either the pre- or post-polishing sequence, indicating that both were correct according to the QV metric. Since it is impossible for both versions of the sequence at a given location to be correct, this 33% of inspected edits likely falls in non-unique sequence, where either the pre- or post-polishing sequence is found elsewhere in the genome, so the QV failed to flag it as false. This proportion may also be caused by noise from sequencing errors in the Illumina reads. In addition, all but 2 of the 42 edits examined were found within long tandem repeats or homopolymers, which are more likely to be non-unique in the genome. We propose that this category accounts for the much lower error rate reported by the QV than by the GIAB analysis within the high confidence regions. Because all of the QV reported errors we examined were corroborated as false by GIAB (and there were only 3 cases where QV indicated an error was fixed while GIAB indicated it was induced), we conclude that the majority of the QV errors within the confidence regions are most likely true errors.

### Comparing different methods for estimating QV

We compared several different methods for calculating QV on our HG005 Hifiasm assembly and found them to be highly variable (**Supplementary Table 6**). Merqury^7^ and Yak^8^ typically report values 2-3 QV points apart (approximately a 2x difference in error rate). We found that using k-mers of size 31 led to more conservative scores than k of 21 (between 2-4 points lower, equivalent to an approximately 50% difference in error rate). These longer k-mers can capture differences in longer repeats, which likely accounts for the higher predicted error rates. We also tested Merqury with a hybrid of HiFi and Illumina k-mers following the approach suggested in McCartney et. al^24^. We found that the hybrid k-mer database led to QV scores around 10 points higher (a predicted order of magnitude lower error rate) than when we used just Illumina. While using hybrid k-mers may remove some Illumina-specific bias, specifically known issues with GC-bias^11,12^, we observed that it introduces too many HiFi-specific sequencing errors to the “truth” k-mer space, and so reduces the apparent error rate by masking true errors. We also experimented with using Element cloudbreak^37^ short read k-mers to calculate QV, and found it predicted double the error rate relative to Illumina (**Supplementary Table 6**). However, we found the same impact of GC bias on the error k-mers calculated by Merqury QV as was described for Illumina. **(Supplementary Figure 4)**. Given that Element data was not available for the HPRC samples, and to be conservative, we report Merqury QV with Illumina k-mers of size 31 for this paper unless otherwise stated.

### Characterizing the context of unchanged error k-mers

After obtaining the error k-mers unchanged by polishing as described in the section “DeepPolisher GQ filter optimization”, we sought to establish how many were located within coverage dropouts. We ran the tool mosdepth with parameter --quantize 0:5:10:150: on the PHARAOH alignment to obtain locations with less than 5x coverage, then bedtools intersect to obtain the error k-mers unchanged by polishing within those regions. To obtain the number that were within simple or low complexity repeats, we ran dipcall with default parameters to align the HG005 raw assembly against CHM13, then the script https://github.com/mobinasri/flagger/blob/main/programs/src/project_blocks_multi_thread.py to project the repeat annotations produced for T2T-CHM13 at https://github.com/marbl/CHM13 to HG005 coordinates. We then used bedtools intersect with the unchanged error k-mers and the projected repeat annotations. To calculate the GC content of all error k-mers we used bedtools nuc. To obtain regions in the HG005 raw assembly with homopolymers >10bp, we used a custom python script which may be found at https://github.com/miramastoras/DeepPolisher_project/blob/main/quantify_error_kmers_HG00.md, and bedtools intersect.

### Gene impact analysis

We annotated the HG002 and HG005 assemblies before and after polishing with the Comparative Annotation Toolkit (CAT2.0)^47^ using the transMap and liftoff modules based on the GENCODE v46 gene annotations for hg38 as the reference. For these annotation sets, we identified the locations of frameshifts by iterating over the coding sequence of every transcript and looking for gaps in the alignment. If the gap length was not a multiple of 3 or if the length was longer than 30bp, the gap was determined to be a frameshift. To identify the number of nonsense mutations that would cause early stop codons in the annotation sets, we iterated through each codon in the coding sequence and looked for an early stop codon before the canonical stop codon at the end of the transcript. To identify the number of missense or nonsynonymous mutations that would cause a change in the amino acid sequence produced, we iterated through each codon, and for codons that were different from the canonical sequence in hg38, checked if the substitution of the nucleotide caused a change in the amino acid produced.

### Element polishing

We obtained 50x element data for HG002 and HG005 from Carroll et. al^37^ and aligned all reads to each polished haplotype assembly separately. We used DeepVariant v1.6.1 with --model_type=WGS to call variants. We selected only homozygous alt PASS variants with genotype quality greater than 7, and applied them to the assembly with bcftools consensus -H2. Code used in this analysis is located at https://github.com/miramastoras/DeepPolisher_manuscript/blob/main/paper_analysis/element_polishing.md.

## Supplementary figures

**Supplementary Figure 1:**
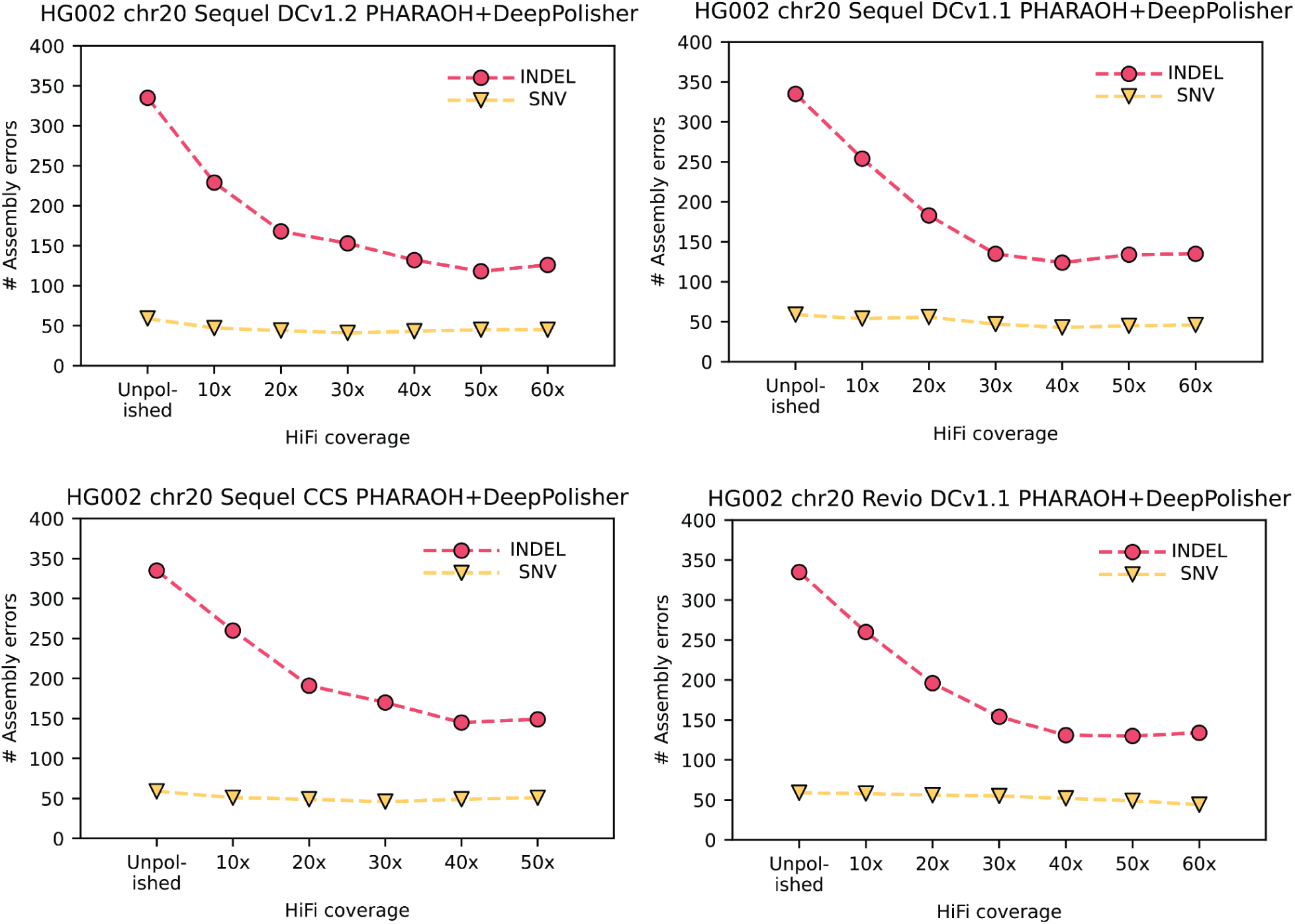
HG002 chr20 GIAB variant calling performance for other HiFi read versions, across different coverages. Total GIAB variant calling (assembly) errors for different HiFi read coverages, with indel errors represented in pink circles and SNV errors in yellow triangles

**Supplementary Figure 2:**
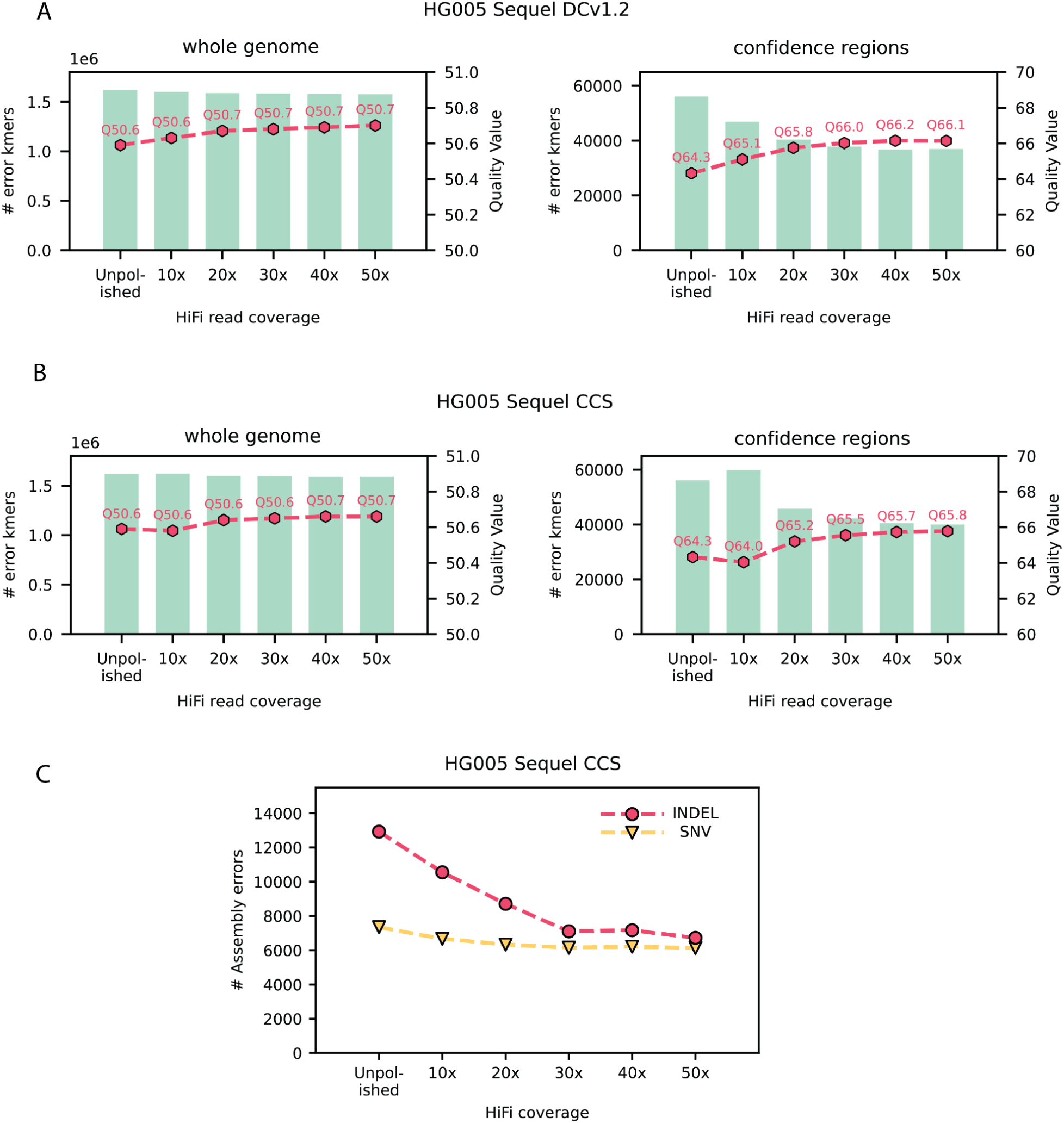
HG005 data performance across multiple coverages and read technologies. **A)** Bar plots represent number of error k-mers (left y axis) for HiFi Sequel DCv1.2 at different read coverages, with the corresponding QV plotted on the right y axis for whole genome (left panel) and for the confidence regions (right panel) **B)** The same for HiFi Sequel CCS data. **C)** Total GIAB variant calling (assembly) errors for different HiFi read coverages, with indel errors represented in pink circles and SNV errors in yellow triangles

**Supplementary Figure 3:**
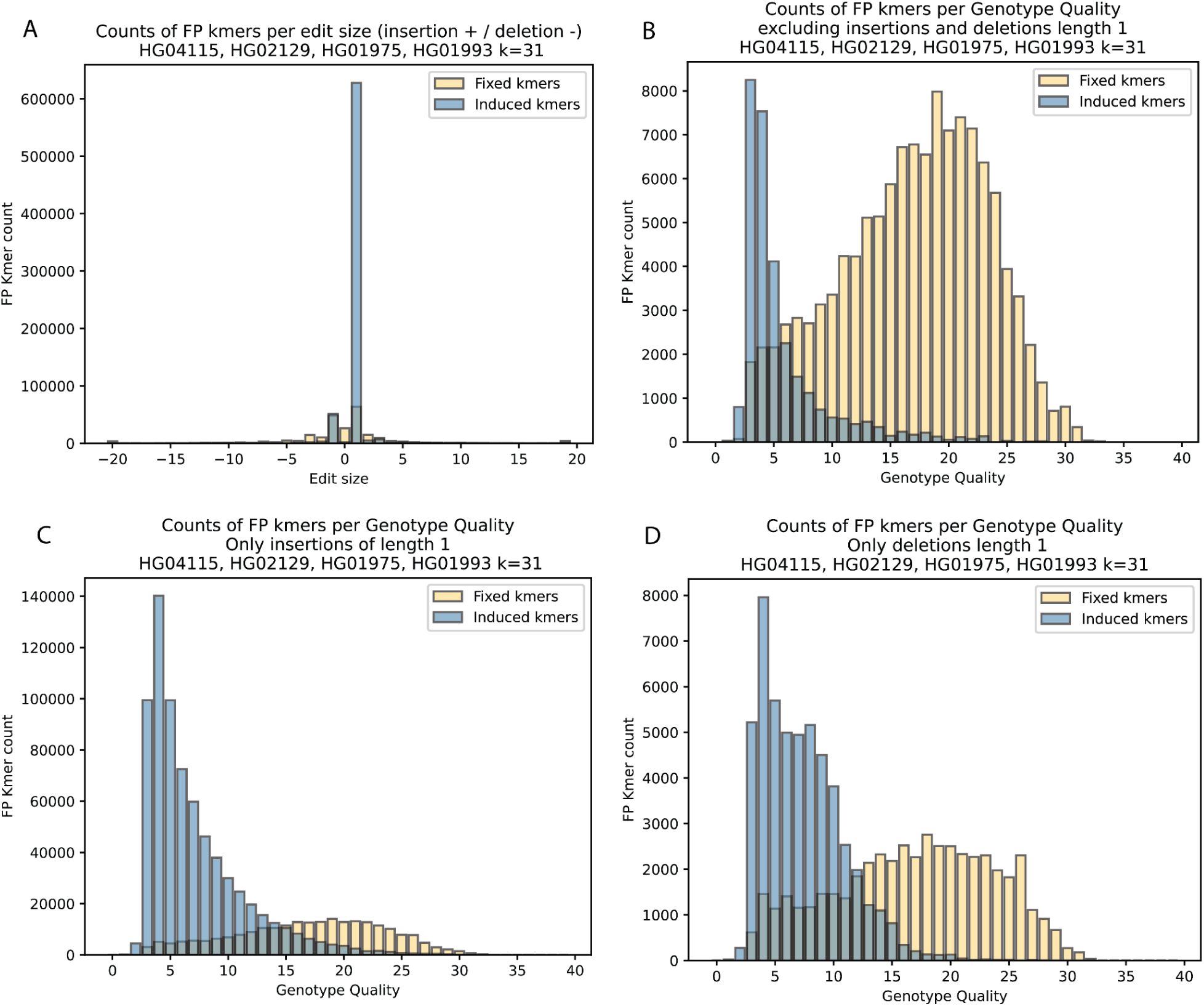
Optimizing GQ filters for DeepPolisher. **A)** Count of error (FP) k-mers per polishing edit size, with error k-mers fixed by polishing edit in yellow and error k-mers induced by polishing edit in blue. **B)** Counts of error (FP) k-mers per genotype quality of polishing edit, excluding insertions and deletions of length 1, and for **C**) the same for only insertions of length 1, and **D)** for deletions of length 1.

**Supplementary Figure 4:**
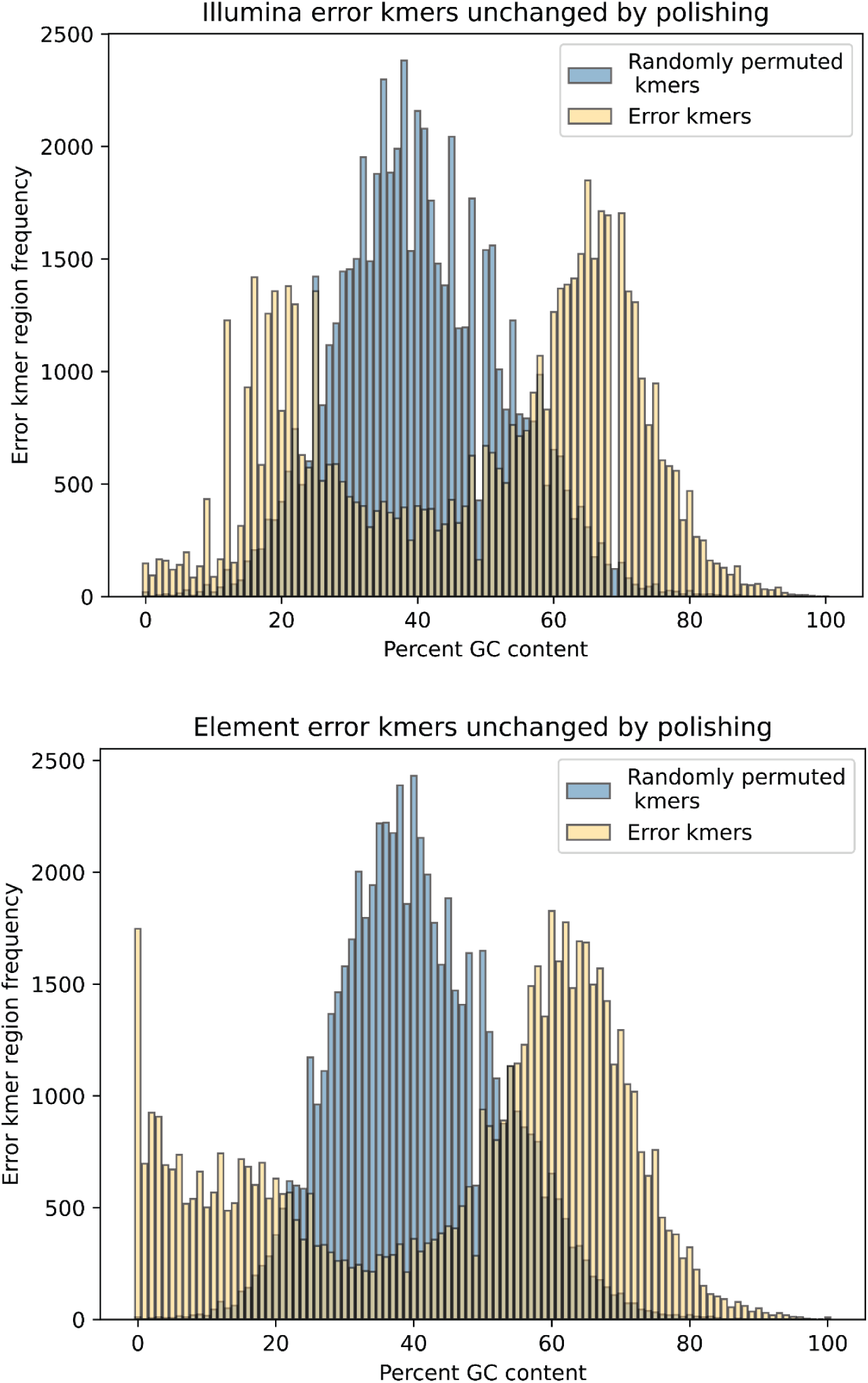
Percent GC content of error k-mers unchanged by polishing and randomly permuted k-mers. Frequency of merged error k-mer regions per varying GC contents. In yellow shows the actual observed error k-mers produced by Merqury, in blue are randomly permuted k-mers of the same size across the genome. Top panel is for Illumina, bottom panel for Element cloudbreak.

